# Neural population dynamics reveals disruption of spinal sensorimotor computations during electrical stimulation of sensory afferents

**DOI:** 10.1101/2021.11.19.469209

**Authors:** Natalija Katic Secerovic, Josep-Maria Balaguer, Oleg Gorskii, Natalia Pavlova, Lucy Liang, Jonathan Ho, Erinn Grigsby, Peter C. Gerszten, Dzhina Karal-ogly, Dmitry Bulgin, Sergei Orlov, Elvira Pirondini, Pavel Musienko, Stanisa Raspopovic, Marco Capogrosso

## Abstract

While neurostimulation technologies are rapidly approaching clinical applications for sensorimotor disorders, the impact of electrical stimulation on network dynamics is still unknown. Given the high degree of shared processing in neural structures, it is critical to understand if neurostimulation affects functions that are related to, but not targeted by the intervention. Here we approached this question by studying the effects of electrical stimulation of cutaneous afferents on unrelated processing of proprioceptive inputs. We recorded intra-spinal neural activity in four monkeys while generating proprioceptive inputs from the radial nerve. We then applied continuous stimulation to the radial nerve cutaneous branch and quantified the impact of the stimulation on spinal processing of proprioceptive inputs via neural population dynamics. Proprioceptive pulses consistently produced neural trajectories that were disrupted by concurrent cutaneous stimulation. This disruption propagated to the somatosensory cortex, suggesting that electrical stimulation can perturb natural information processing across the neural axis.

## Introduction

Decades of animal and human studies have shown that neurostimulation technologies can restore some level of neurological function in patients with sensorimotor deficits(*1–10*). These novel technologies produce immediate assistive effects, achieving a controlled restoration of multifaceted behavioral processes(*11*). For instance, in humans peripheral neuroprostheses successfully restore touch sensations(*12–20*), and spinal cord stimulation enables the recovery of voluntary motor control(*3–5*). While these remarkable results are fueling the translation of these technologies in clinical settings, the understanding of the short- and long-term effects of injecting electrical current into existing neural dynamics is still entirely unknown. In fact, virtually all these interventions suffer from a latent, yet critical caveat: the input delivered to the neural circuits is artificially generated, being widely different from naturally-generated neural activity.

Indeed, electrical stimulation produces synchronized volleys of neural activity in all recruited axons (or cells), rather than the asynchronous bursts of inputs that govern natural neural activity(*21, 22*). What is the consequence of this stark difference with respect to neural function? Recently, some studies demonstrated that electrical stimulation actually triggers side effects at the neural level, which were initially unnoticed. For example, new data from epidural spinal cord stimulation for spinal cord injury showed that continuous stimulation of recruited sensory afferents produces a disruption of proprioceptive percepts at stimulation parameters commonly employed in clinical trials(*6*). Similarly, the inability to elicit robust proprioceptive percepts(*23*) is striking in the application of electrical stimulation of the peripheral nerves for the restoration of somato-sensations.

In fact, large-diameter proprioceptive afferents should have the lowest threshold for electrical stimulation. Therefore, these afferents should be the easiest sensory afferents to recruit with neural interfaces(*24–28*). However, because of anatomical and geometrical constraints, in practice electrical neurostimulation leads to the activation of mixed diameter fiber distributions and, consequently, different sensory modalities (*8, 25, 29*). Therefore, cutaneous afferents are recruited concurrently, along with larger diameter afferents(*30*). These fibers converge on interneurons in the spinal cord where they undergo the first layer of sensory processing, representing a highly shared sensory network node.

It is conceivable that artificially-generated patterns of mixed neural activity hinder some of the computations of these shared network nodes, thus impairing natural circuit processing, hence, perception (**Fig 1**). In order to demonstrate this conjecture, we need tools that allow us to visualize and identify a direct measure of neural computation processes(*31*). Analysis of population neural dynamics using neural manifolds is commonly employed to study computational objects that process information in the cortex(*32–35*) and, more recently, in the spinal cord. Indeed, one study showed that intraspinal population responses contain simple structures that enable the examination of complex processes such as walking(*36*).

**Fig. 1.**
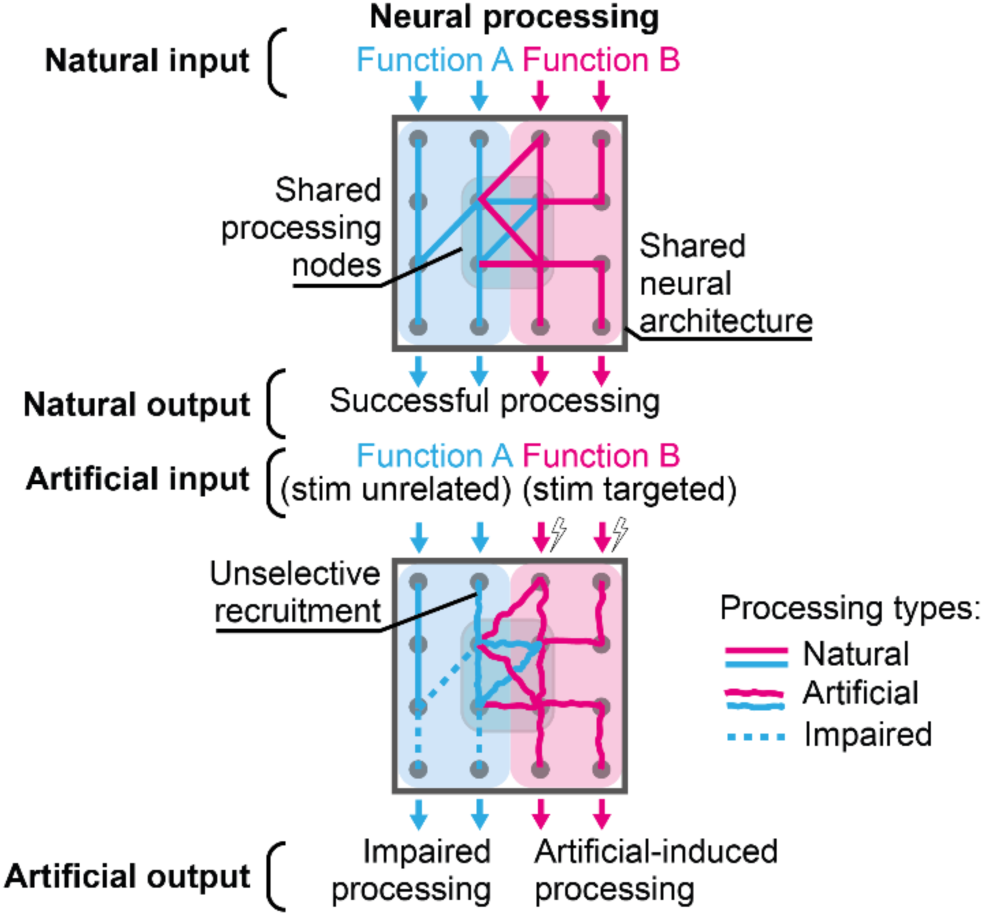
Electrical stimulation disrupts computations of ongoing network processes. Neural networks (cyan, magenta shaded areas) produce a desired neural function (Function A, B, respectively) within a highly shared neural architecture. These networks may share processing layers (or nodes) to process input information. Top, naturally-generated neural activity of unrelated neural functions successfully processes ongoing information input throughout the shared neural architecture. Bottom, artificially-generated neural activity targeted to restore Function B (magenta) artificially processes information input, while impairing information processing from an unrelated neural function (Function A, cyan). Specifically, artificially-induced processing in the shared processing nodes concurrently hinders computations of unrelated ongoing processing of Function B, which may also be unselectively recruited by the electrical stimulation.

Therefore, here we employed neural population analysis of intraspinal neural dynamics to 1) visualize neural computations underlying the processing of brief proprioceptive percepts elicited by single short pulses of electrical stimulation and 2) study how these computations were altered when concurrent electrical stimulation was delivered to sensory afferents from a different nerve. This experimental design offered a simplified version of the more general problem of the stimulation effects on unrelated neural functions, thus allowing us to execute casual manipulation and quantification of neural variables.

Therefore, we designed a series of electrophysiology experiments in anesthetized monkeys, who share distinguishable projections with the human nervous system distinct from all other animals(*37, 38*). We recorded and analyzed artificially evoked proprioceptive neural signals both in the cervical spinal cord and somatosensory cortex. Specifically, we induced proprioceptive input in the hand and forearm by cuff electrode stimulation of the muscle branch of the radial nerve, which does not contain cutaneous afferents(*39, 40*). Then, we studied how concurrent stimulation of somatosensory afferents in the cutaneous branch of the radial nerve impacted the spinal and cortical proprioceptive responses. Using neural population analysis, we examined dorso-ventral intra-spinal spiking activity in response to muscle nerve stimulation pulses and performed dimensionality reduction to observe the spinal neural trajectories. Concurrent stimulation of the cutaneous afferents disrupted these neural trajectories, suggesting a significant degradation of proprioceptive information processing in the spinal cord. Changes in proprioceptive information appeared as reduced cortical responses in the somatosensory cortex. Our results show that intraspinal neural population dynamics can capture the processing of sensorimotor information in spinal networks and its disruption of this information processing during artificial electrical stimulation.

## Results

### Simultaneous brain and spinal neural recordings during electrical nerve stimulation of multiple sensory modalities

We designed a unique experimental setup in non-human primates as a proxy to understand how artificial inputs can influence neural network function in a controlled fashion. Specifically, we examined how stimulation of cutaneous afferents affects spinal network processing of proprioceptive pulses. The radial nerve, carrying sensory signals from the dorsal part of the forearm and hand, splits in proximity of the elbow into a pure-muscle and a pure-cutaneous branch (i.e., the deep and superficial branches of the radial nerve(*40*), respectively) offering the opportunity to provide modality-selective sensory stimuli. We implanted cuff electrodes on these two branches to elicit either proprioceptive or cutaneous inputs via electrical stimulation (**Fig. 2a**).

**Fig. 2.**
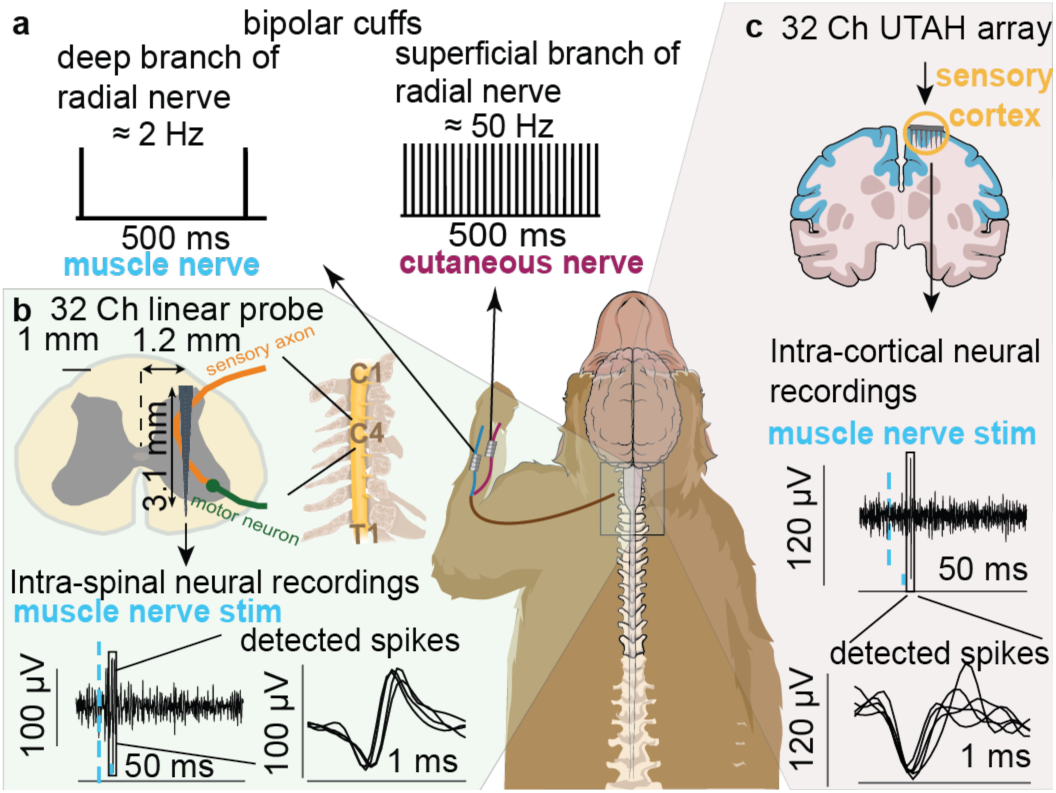
Experimental setup: Schematic illustration of experiments. a) Stimulation: we implanted two nerve cuffs for stimulation on the superficial branch (cutaneous nerve) and the deep branch (muscle nerve) of the radial nerve. We stimulated the muscle nerve at ~2 Hz, exclusively, or concurrently with ~50 Hz stimulation of the cutaneous nerve branch. b) We recorded neural activity with a 32-channel dorso-ventral linear probe implanted in the gray matter of the spinal segment C5. Typical intra-spinal neural responses induced by stimulation of the muscle nerve. Zoom insets show examples of detected spike waveforms, e.g., singleunit responses to proprioceptive pulses. c) We recorded neural activity with a 32-channel multi-electrode array in the somatosensory cortex and provided intra-cortical neural responses, similar as in b.

We artificially provided brief proprioceptive pulses by stimulating the muscle branch of the radial nerve with single electrical pulses (~2 Hz) below motor threshold. To assess the influence of artificial cutaneous input on the induced proprioceptive input, we provided cutaneous stimulation as continuous ~50 Hz pulses, a typical stimulation frequency used in human studies. Threshold (Thr) was defined as an amplitude that clearly evoked potentials in the spinal cord in response to low-frequency stimulation. We tested two conditions: stimulating the nerve at a low (0.9 x Thr) or high amplitude (1.1 x Thr). Stimulation amplitude corresponds to the amount of artificially recruited fibers. To study the transmission of artificially induced proprioceptive percepts from the periphery to the cerebral cortex, we recorded the Macaque monkeys’ intra-spinal neural signals from a dorso-ventral 32-channel linear probe implanted in the gray matter of the spinal cord C5 segment (**Fig. 2b**). Furthermore, we extracted intra-cortical neural signals (**Fig. 2c**) using a 32-channel UTAH array placed in the somatosensory cortex (Area S1/S2, **Fig. S1**).

In summary, we recorded neural signals in the spinal cord and the somatosensory cortex of three anesthetized Macaca Fascicularis (MK1, MK2, MK4) and one Macaca Mulatta (MK3) monkey while stimulating only proprioceptive, or concurrently proprioceptive and cutaneous afferents.

### Proprioceptive inputs elicit robust trajectories in the spinal neural manifold

We explored the effect of brief pulses of artificially-generated proprioceptive inputs on the intraspinal neural population dynamics. Because proprioceptive signals enter the spinal cord from the dorsal aspect and project towards medial and ventral laminae(*41*), we performed neural population analysis of the multiunit spiking data from all the channels of our linear probe (**Fig. 3a**) in response to 2 Hz muscle nerve stimulation. Specifically, we applied dimensionality reduction to unveil the latent properties of the spinal neural processing via principal component analysis (PCA). PCA identified three neural modes that sufficed to explain 54-65% of the variance of the spikes counts of multiunit threshold crossings recorded by the spinal probe for ~350 ms following each proprioceptive stimulus pulse. We then sought whether the neural manifold defined by these neural modes contained simple computational objects (e.g., clear neural trajectories that captured the changes of time-varying spikes, **Fig. 3b, c**).

**Fig. 3.**
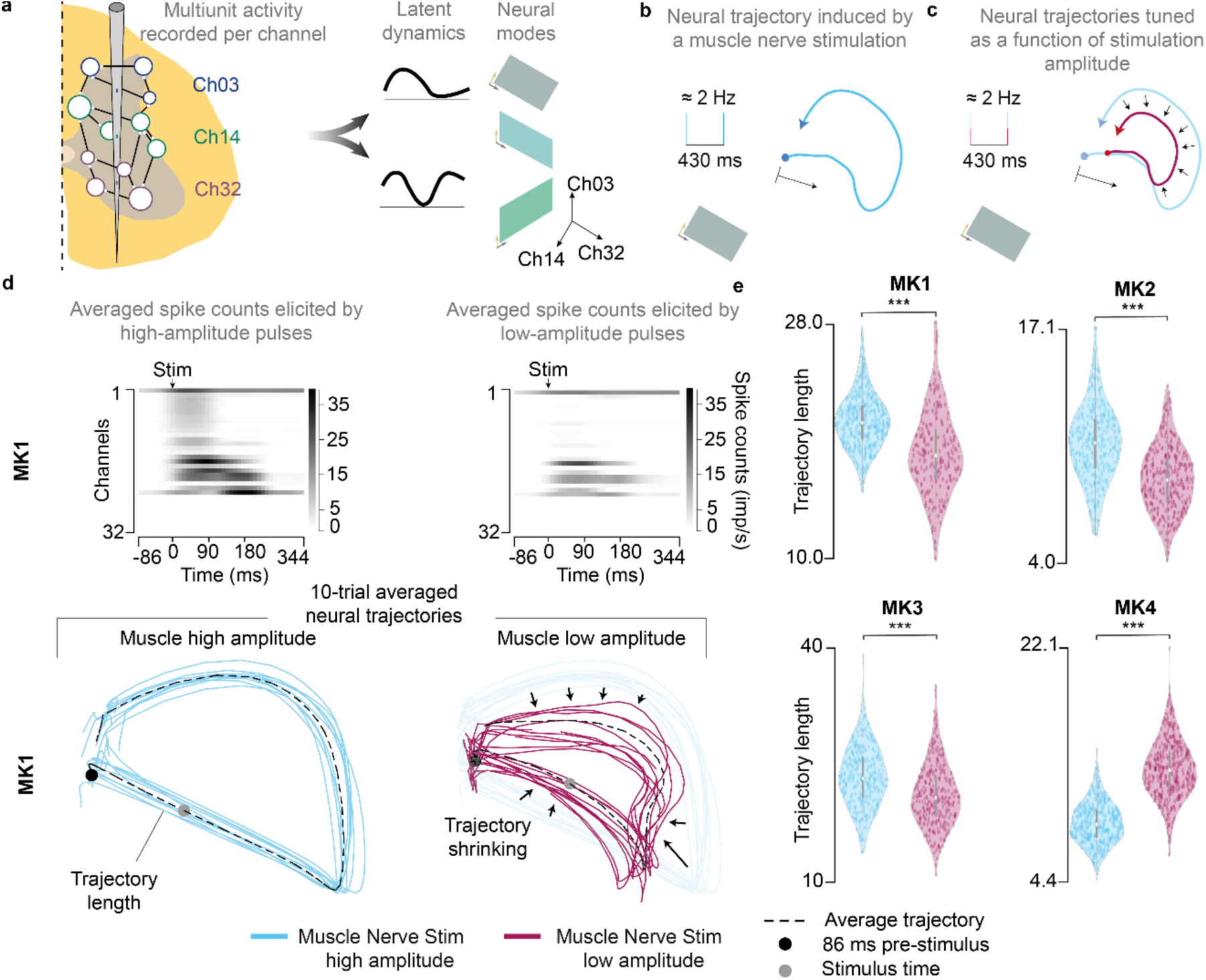
Intra-spinal neural population analysis. a) Latent dynamics and neural modes obtained from the multiunit recorded per channel. Left, a sketch of the dorso-ventral linear probe that recorded the activity of the spinal multiunit neural networks (each circle represents a recorded unit). Each color represents the neural activity recorded by each channel. Right, dimensionality reduction technique identifies the neural modes that define the low-dimensional spaces. In these subspaces, the neural activity followed precise dynamics. We hypothesized that b) a muscle nerve stimulation pulse elicits neural trajectories and that c) these neural trajectories shrink as a function of the stimulation amplitude. d) Top, averaged multiunit spike counts across all 32 channels, sorted by the highest spiking activity after the muscle nerve stimulation, for MK1. Bottom, resultant 10-trial averaged neural trajectories elicited by muscle nerve stimulation for MK1. This is plotted both at a high and low stimulation amplitude to appreciate the phenomenon of trajectory shrinking. e) Statistical quantification of the trajectory length for all monkeys for high and low stimulation amplitude of the muscle nerve (***p<0.001; **p<0.01; *p<0.05; Kruskal-Wallis test with 380 and 231 points for high and low amplitude, respectively, for MK1; 353 and 351 points for high and low amplitude, respectively, for MK2; 353 and 343 points, respectively, for MK3; 391 and 394 points, respectively, for MK4). Violin plots: each dot corresponds to the computed trajectory length for a trial, forming a Gaussian distribution of trajectory lengths. The central mark represented as a white dot indicates the median, and the gray line indicates the 25th and 75th percentiles. The whiskers extend to the most extreme data points not considered outliers. Trial corresponds to a stimulation pulse.

In the spinal manifold, the multiunit spike counts elicited very consistent dynamics after each stimulation pulse in the form of closed trajectories that were qualitatively similar in all monkeys **(Fig. 3d)**. Because averaged spiking responses initiated and terminated with baseline activity (i.e., no stimulation), the neural dynamics were represented by closed neural trajectories. Given the robustness and reproducibility of these trajectories, we hypothesized that estimated trajectory lengths could be used as a proxy to measure the amount of proprioceptive information processed within the recorded site. The logical consequence of this interpretation is that the length of the trajectories could be proportional to the amount of proprioceptive input processed.

Since the stimulation amplitude controls the number of recruited afferents, we tested this assumption by computing the neural trajectories induced by proprioceptive inputs both at high and low stimulation amplitudes (i.e., more or less recruited afferents, respectively). As expected, we found that muscle nerve stimulation at a higher amplitude elicited longer trajectories and vice versa **(Fig. 3d)**. This observation was consistent in MK1 (relative mean difference, +14.17%), MK2 (+24.05%) and MK3 (+44.21%, **Fig. 3e**), but not in MK4 (−33.76%), probably due to the higher variability in the overall trajectories for this monkey.

In summary, we showed that population analysis of a dorso-ventral linear probe in the spinal cord shows highly robust and reproducible trajectories in the neural manifold in response to artificial proprioceptive pulses. We proposed to quantify the length of this trajectory as a means to assess the amount of proprioceptive information processed in the spinal cord.

### Continuous electrical stimulation of the cutaneous nerve disrupts intra-spinal proprioceptive neural trajectories

We next evaluated the impact of concurrent artificial cutaneous input on proprioceptive information processing. We projected on the neural manifold neural trajectories elicited by the stimulation of the proprioceptive branch. All four monkeys exhibited robust trajectories in response to proprioceptive inputs and, in all four monkeys, concurrent stimulation of cutaneous afferents significantly reduced the trajectory lengths (**Fig. 4a**) or even completely disrupted their dynamics (**Fig. S4**), albeit with different effect sizes. MK1 (relative mean difference, −66.03%), MK3 (−27.47%) and MK4 (−44.91%) exhibited the largest disruption, while MK2 (−5.89%) was significantly disrupted but to a lower effect size.

**Fig. 4.**
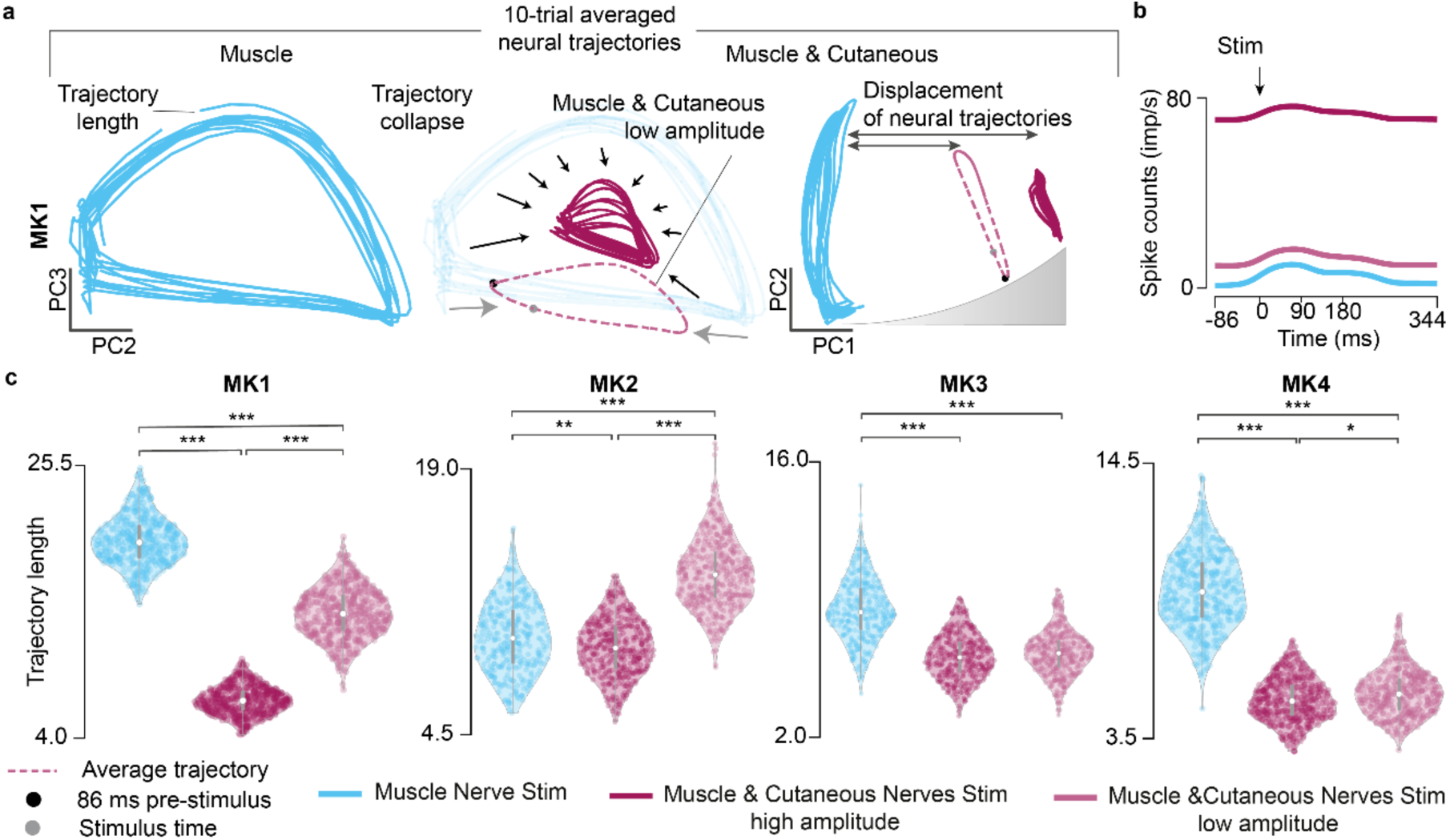
Neural trajectory lengths. a) Comparison of the neural trajectories induced by muscle nerve stimulation and concurrent cutaneous stimulation across PC2-PC3 vs PC1-PC2. Gray dashed lines indicate average trajectory for muscle and cutaneous nerves stimulation at a low amplitude. b) Averaged spike counts across all trials and all channels for each stimulation condition for MK1. c) Statistical analysis of the trajectory lengths for each stimulation condition. Violin plots: each dot corresponds to the computed trajectory length for a trial, forming a Gaussian distribution of trajectory lengths. The central mark represented as a white dot indicates the median, and the gray line indicates the 25th and 75th percentiles. The whiskers extend to the most extreme data points not considered outliers. Trial corresponds to a stimulation pulse. (***p<0.001; **p<0.01; *p<0.05; Kruskal-Wallis test with 381, 470 and 453 points for muscle nerve stimulation, concurrent cutaneous stimulation at high amplitude and low amplitude, respectively, for MK1; 369, 410 and 411 points, respectively, for MK2; 353, 376 and 397 points, respectively, for MK3; 392, 380 and 371 points, respectively, for MK4).

To validate this result, we repeated the same experiment using lower amplitudes for the stimulation of the cutaneous afferents. Cutaneous stimulation at a low amplitude yielded less disruption (i.e., longer proprioceptive trajectory lengths) than at a high stimulation amplitude (**Fig. 4c**), suggesting that the amount of neural computation disrupted is inversely proportional to stimulation intensity of cutaneous afferents. This observed trajectory disruption is particularly interesting considering that, during concurrent proprioceptive and cutaneous stimulation, the spinal cord received significantly more artificial input. Indeed, concurrent stimulation of the cutaneous afferents significantly increased overall spike counts in the recorded spinal circuitries (+5191.68% for MK1, +234.71% for MK2, +68.37% for MK3, +754.94% for MK4, **Fig. 4b, S3**). However, this increase was not captured by the neural trajectories, which strengthens the case that those trajectory lengths mainly represent proprioceptive information processing.

Additionally, we found that the disruption of information processing was captured in principal component (PC) 2 and PC3, where neural trajectories shrunk as a function of stimulation intensity. Moreover, PC1 depicted the displacement of these neural trajectories caused by the amount of concurrent cutaneous input (**Fig. 4a, Fig. S3**). In other words, when the spinal cord received inputs induced by concurrent muscle and cutaneous nerves stimulation, the neural trajectories were displaced across PC1, away from the proprioceptive neural trajectories. This displacement was proportional to the stimulation intensity and, in turn, to the computed spiking activity in the spinal cord (**Fig. S3**). In particular, MK1 (+94.52%) and MK2 (+45.99%) produced longer proprioceptive trajectory lengths than MK3 (+3.49%) and MK4 (+5.22%) during concurrent cutaneous stimulation at a low amplitude. Indeed, the overall spike counts were very similar in MK3 and MK4 both at low and high stimulation amplitudes (**Fig. S3**) (relative mean difference from concurrent high to low amplitude, −84.44% for MK1, −59.82% for MK2, −1.69% for MK3, −8.40% for MK4), thereby eliciting similar neural trajectory lengths. These results infer that the main PCs clearly captured the amount of proprioceptive processed information as a function of concurrent stimulation amplitude. Indeed, neural trajectories that were further displaced across PC1 resulted in shorter neural trajectory lengths in PC2-PC3 (during concurrent high stimulation amplitude), whereas those that remained closer to the proprioceptive neural trajectories in PC1 were less disrupted in PC2-PC3 (during concurrent low stimulation amplitude, **Fig. 4a**).

In summary, we showed that concurrent stimulation of the cutaneous nerve significantly suppressed proprioceptive neural trajectory lengths, suggesting that concurrent artificial recruitment of cutaneous afferents hinders the processing of proprioceptive inputs in the spinal cord.

### Cutaneous electrical stimulation reduced proprioceptive afferent volleys, spinal cord grey matter field potentials and multiunit responses

To validate our findings, we looked for correlates using classical electrophysiology measures. We first inspected stimulation triggered average field potentials from the grey matter of the spinal cord, defined as the mean neural response across each single muscle nerve branch stimulation pulse (**Fig. 5a**). Afferent volleys were detected at a latency between 3-4 ms after each proprioceptive pulse (**Fig. S5 and Fig 5a**). Continuous electrical stimulation of the cutaneous nerve reduced the peak-to-peak amplitude of these proprioceptive volleys in all four monkeys **(Fig. S5)** and the reduction was proportional to stimulation intensity (muscle nerve stimulation vs muscle & cutaneous nerve stimulation high amplitude, mean values difference: MK1: −9%, MK2: −47%, MK3: −25%, MK4: −14%; muscle nerve stimulation vs muscle & cutaneous nerve stimulation low amplitude: MK1: −8%, MK2: −40%, MK3: −22%, MK4: −1%). Since volleys represent sensory inputs, these results suggest that part of the disruption that we observed in the neural trajectories may be a consequence of reduced proprioceptive inputs in the spinal cord.

**Fig. 5.**
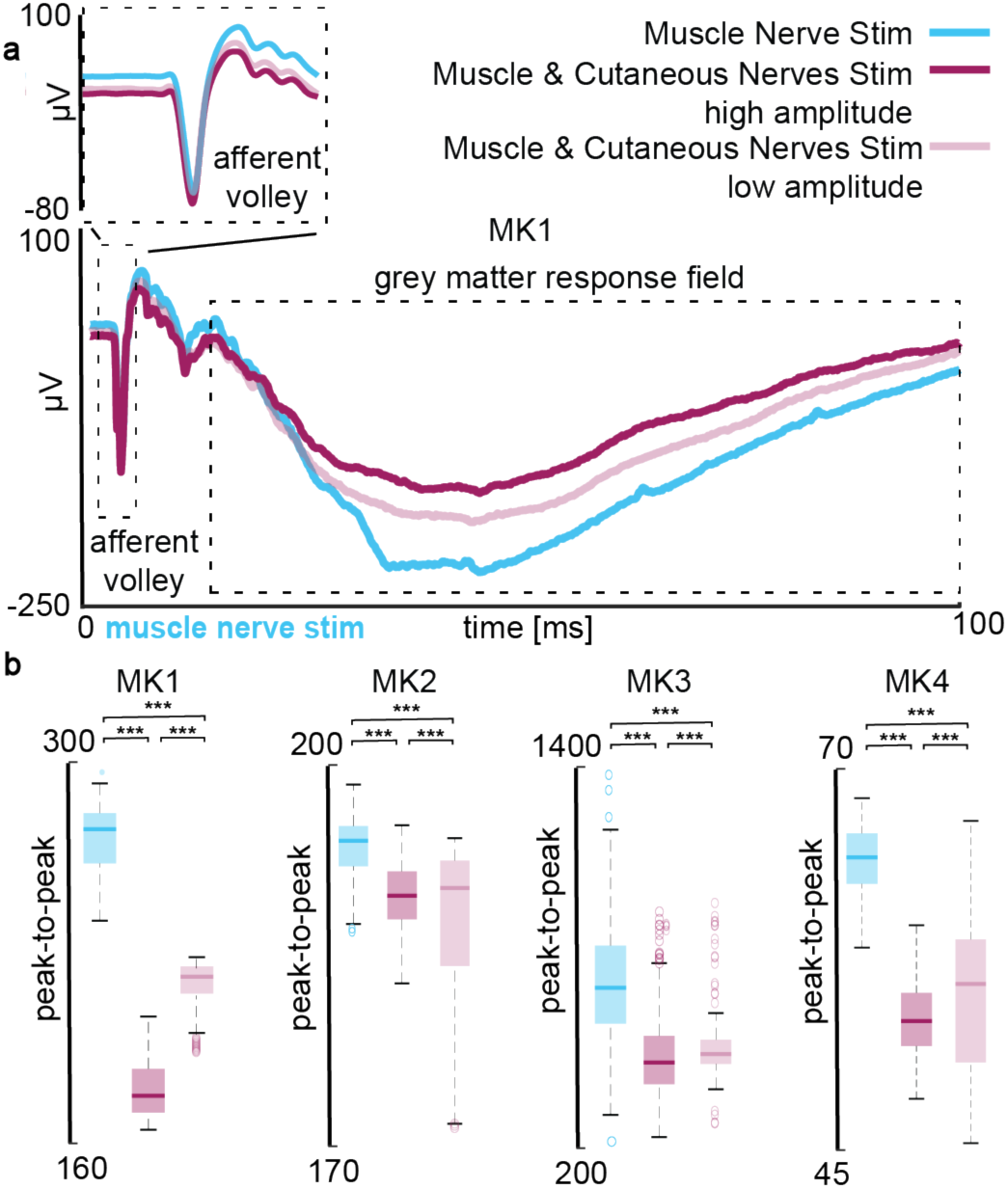
Peak-to-peak amplitude suppression of spinal cord grey matter response fields. a) MK1 triggered-average signal showing afferent volley, and grey matter response fields resulting from muscle nerve stimulation (cyan), with concurrent cutaneous nerve stimulation (magenta; high stimulation amplitude – solid color; low stimulation amplitude – semi-transparent). b) Peak-to-peak amplitude of grey matter response field in four monkeys, dorsal channels examples. Color coding the same as in a. We compared peak-to-peak amplitude values over two conditions with one-way ANOVA with 300 points, where each point represents the peak-to-peak amplitude as a response to a single stimulus pulse. Boxplots: The central mark indicates the median, and the bottom and top edges of the box indicate the 25th and 75th percentiles, respectively. The whiskers extend to the most extreme data points not considered outliers, and the outliers are plotted individually using the ‘o’ symbol. Asterisks: ***p<0.001.

Additionally, grey matter response fields following each proprioceptive volley were also substantially suppressed during electrical stimulation. Peak-to-peak amplitude values of the fields were significantly reduced to a much larger extent than the volleys. Again, the suppression correlated to stimulation intensity: high amplitude of cutaneous stimulation resulted in greater suppression of afferent volleys and grey matter response fields peak to peak values than at a low stimulation amplitude (muscle nerve stimulation vs muscle & cutaneous nerve stimulation high amplitude, mean values difference: MK1: −83%, MK2: −18%, MK3: −46%, MK4: −56%; muscle nerve stimulation vs muscle & cutaneous nerve stimulation low amplitude: MK1: −48%, MK2: −15%, MK3: −42%, MK4: −43%, **Fig. 5b**). The same trend was found in all 4 monkeys, suggesting that a significant component of the trajectory disruption may be related to reduced grey matter responses to proprioceptive volleys and not only to a simple reduction of proprioceptive inputs.

Finally, we investigated whether changes in the population neural dynamics and grey matter field potentials could be reflected in changes of single neuron spiking activity. We utilized multiunit threshold crossing analysis (**Fig. 6a**) and identified channels in which a clear response to proprioceptive pulses was visible. In this multiunit analysis, the peak of neural activity after proprioceptive stimuli occurred at approximately 3 – 4 ms after each proprioceptive stimulation pulse. We present the neural responses of units that were activated by proprioceptive inputs. When continuous stimulation of the cutaneous nerve was overlapped with muscle nerve stimulation, we observed a reduction in these responses in all four monkeys, both in the dorsal and ventral horn of the spinal cord (**Fig 6b**).

**Fig. 6.**
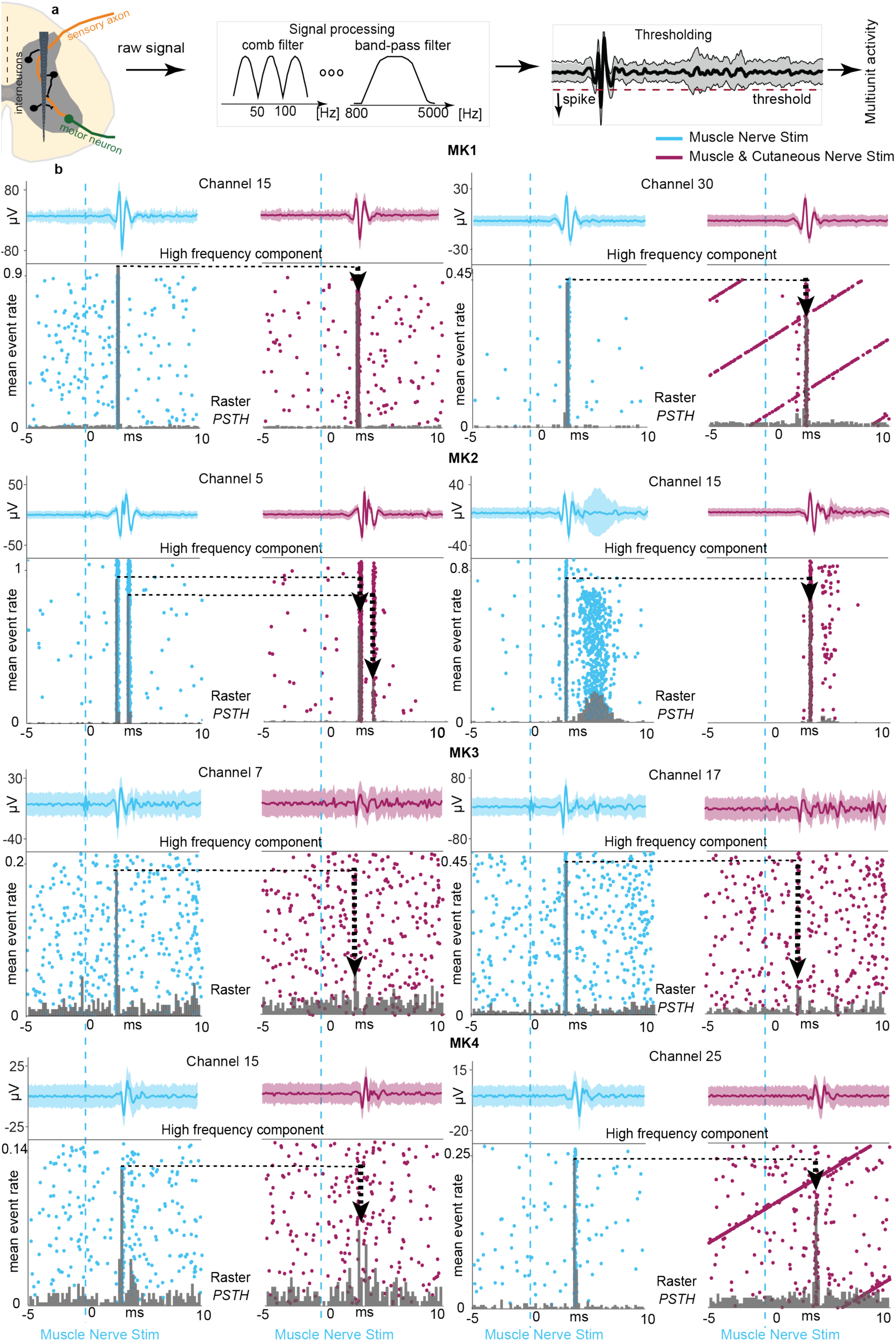
Multiunit activity. a) We filtered the signal to extract the spiking component and detected the neural action potentials using the thresholding algorithm (see methods). b) Examples of multiunit activity in two different channels (one in dorsal, one in ventral region) for each of the four monkeys. Single muscle nerve stimulation (cyan, left) and concurrent muscle and cutaneous nerve stimulation at a high amplitude (magenta, right). Dashed cyan line represents the muscle nerve stimulation pulse. Neural activity is presented and quantified with raster plots and peri-stimulus time histograms (PSTHs). Each row of the raster plots represents the response to a single muscle nerve stimulation pulse, while each dot corresponds to an action potential. Mean event rate is defined as an average number of spikes within a time frame of one bin (0.2 ms) across all single pulses of muscle nerve stimulation. Black lines highlight the PSTH bins that are reduced. Black arrows indicate the decreased mean event rate values of PSTH and their lengths correspond to the amount of reduction. Diagonal lines correspond to the units whose frequency is in line with frequency of stimulation.

In summary, we found that concurrent cutaneous nerve stimulation reduced peak-to-peak amplitude of afferent volleys, grey matter response fields and multiunit responses to proprioceptive stimuli. These results suggest that proprioceptive information processing may be disrupted by reducing both sensory input in the spinal cord as well as grey matter network computations.

### Reduction of proprioceptive processing impacts somatosensory cortex

We showed that concurrent stimulation of cutaneous afferents suppresses proprioception information processing in the spinal cord and correlates to classic electrophysiology measures. We then hypothesized that this suppression in the spinal cord limits the amount of information transmitted upstream to the brain, which could impact conscious perception of proprioception.

To test this hypothesis, we analyzed intra-cortical neural signals extracted from area 2 of the somatosensory cortex in all four monkeys. We found cortical evoked potentials with a latency of around 22-25 ms, which is consistent with the longer distance between the cortex and the peripheral nerves and it has also been reported in similar experiments(*42*). Peak-to-peak analysis of the signal amplitude indicated similar results as in the spinal cord. We observed a reduction of proprioceptive evoked potentials during concurrent high amplitude stimulation of the cutaneous nerve in all monkeys (**Fig. 7**).

**Fig. 7.**
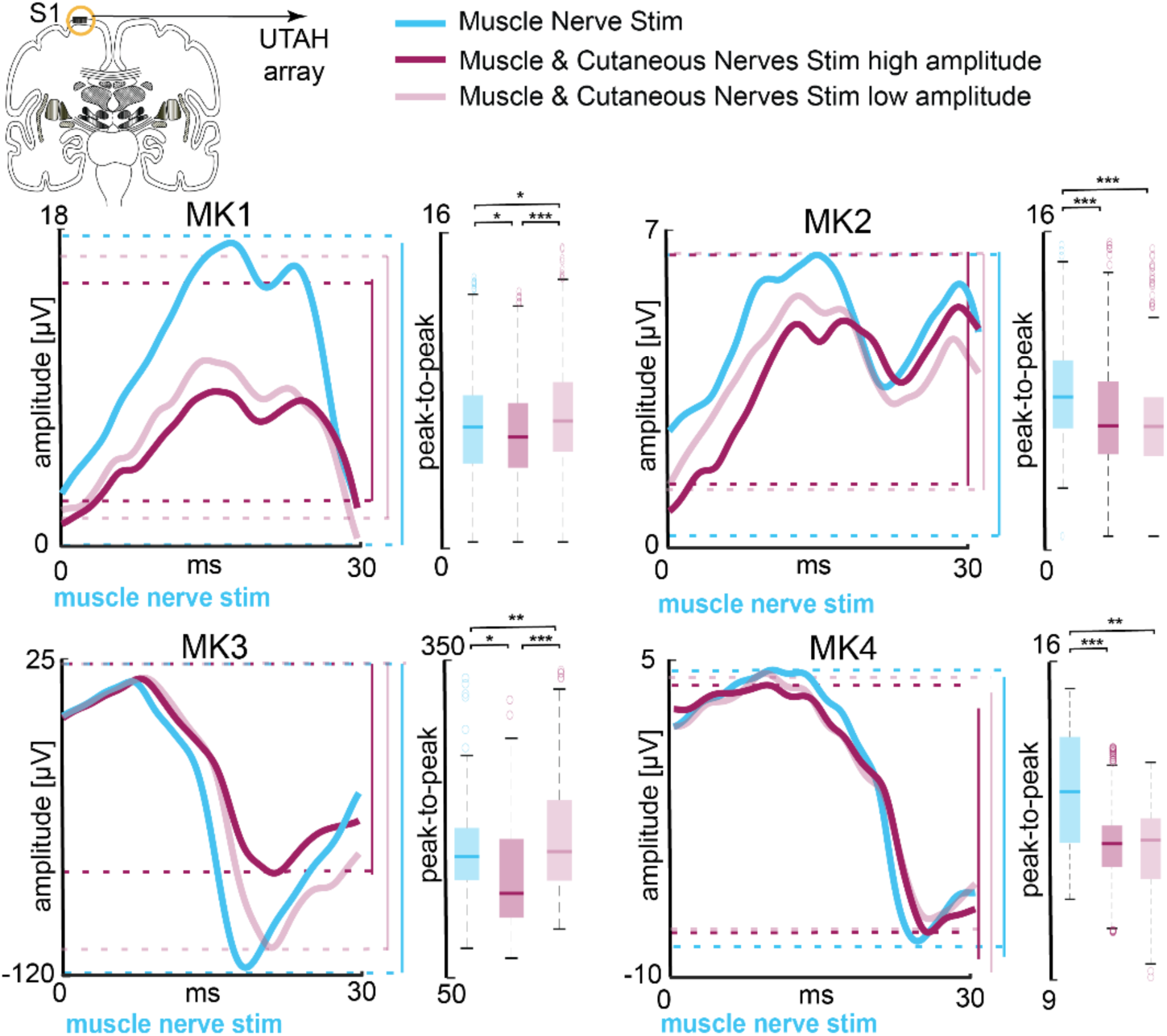
Peak-to-peak amplitudes of muscle nerve stimulation evoked potentials are suppressed in the somatosensory cortex. Somatosensory cortex evoked potentials in four monkeys. Examples of signals recorded as a response to muscle nerve stimulation, with concurrent cutaneous nerve stimulation (magenta; high stimulation amplitude – solid color; low stimulation amplitude – semi-transparent) or without it (cyan). Evoked potentials appeared with a latency between 22-25 ms. Signals are given as an example of a single channel in the somatosensory cortex and are averaged across all muscle nerve stimulation pulses. We compared peak-to-peak amplitude values of the signal over two conditions with one-way ANOVA with 300 points, where each point represents the peak-to-peak amplitude as a response to a single stimulus pulse. Boxplots: The central mark indicates the median, and the bottom and top edges of the box indicate the 25th and 75th percentiles, respectively. The whiskers extend to the most extreme data points not considered outliers, and the outliers are plotted individually using the ‘o’ symbol. Asterisks: ***p<0.001; **p<0.01; *p<0.05.

Observed suppression was detected in most of the channels in the array. Moreover, when we stimulated the cutaneous nerve at a low amplitude, peak-to-peak values of the signal increased (muscle nerve stimulation vs muscle & cutaneous nerve stimulation high amplitude, mean values difference: MK1: −8%, MK2: −19%, MK3: −30%, MK4: −29%; muscle nerve stimulation vs muscle & cutaneous nerve stimulation low amplitude: MK1: +2%, MK2: −18%, MK3: +3%, MK4: −26%).

Surprisingly, when we inspected the spiking activity extracted from multiunits in the cortex, the spike counts induced by concurrent cutaneous nerve stimulation at a low amplitude were similar, or even greater, than those obtained at a high amplitude (relative mean difference from concurrent high to low amplitude, +53.69% for MK1, −7.65% for MK2, +3.14% for MK3, −6.40% for MK4, **Fig. 8**). This is markedly different from what we observed in the spinal cord (**Fig. S2, S3**). Indeed, we expected greater spiking activity consistently associated with higher stimulation amplitudes and not the opposite. In fact, this discrepancy seemed to reflect the spinal proprioceptive information processing, where concurrent cutaneous stimulation at a low amplitude yielded longer neural trajectory lengths.

**Figure 8.**
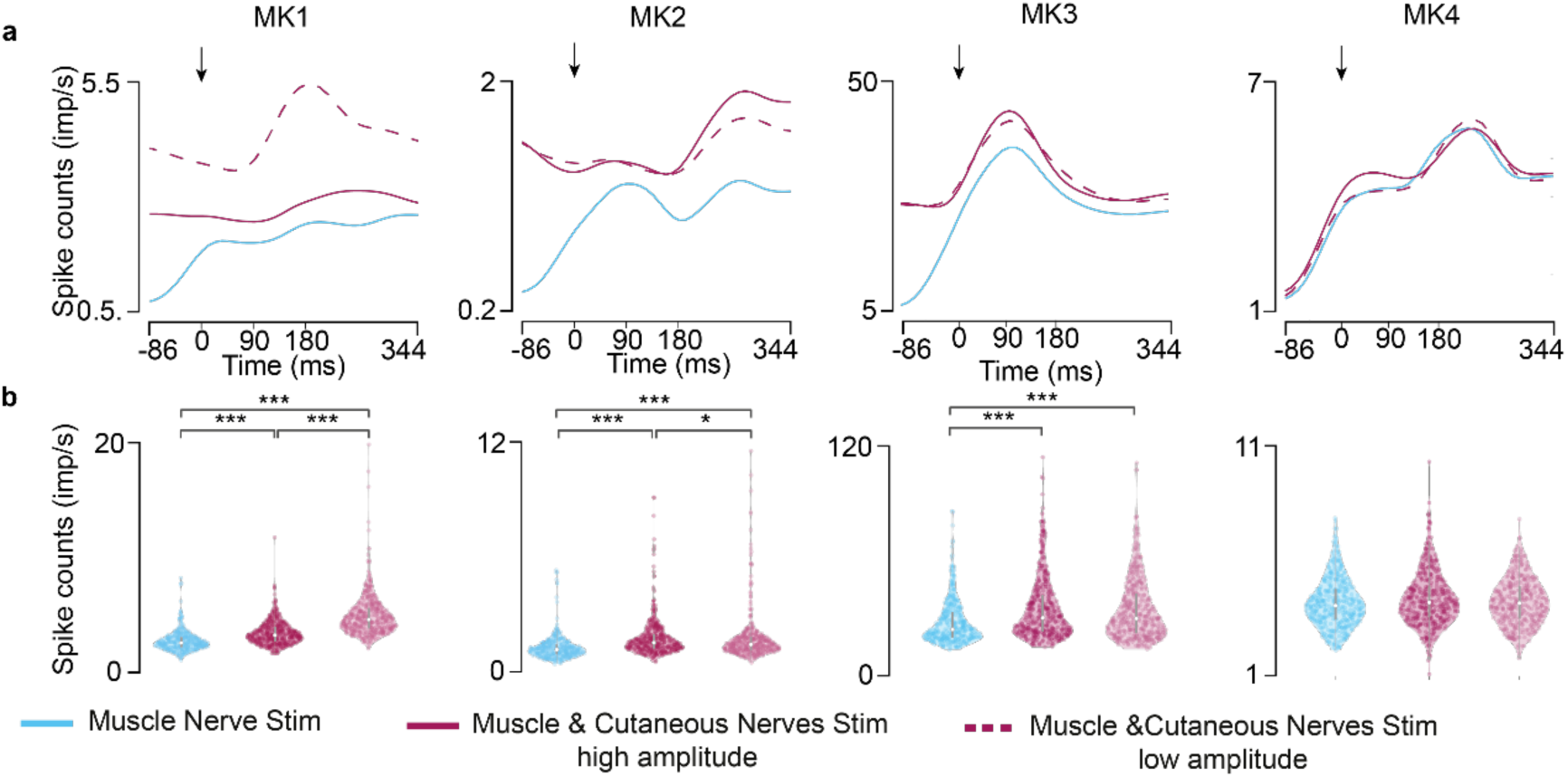
Cortical spiking activity. a) Spike counts were averaged across all trials and all channels for each stimulation condition for all monkeys. b) Statistical analysis of the spiking activity for each stimulation condition (***p<0.001; **p<0.01; *p<0.05; Kruskal-Wallis test with 392, 474 and 460 points for muscle nerve stimulation, concurrent cutaneous stimulation at high amplitude and low amplitude, respectively, for MK1; 360, 389 and 392 points, respectively, for MK2; 335, 386 and 390 points, respectively, for MK3; 391, 383 and 375 points, respectively, for MK3). Violin plots: each dot corresponds to the computed trajectory length for a trial, forming a Gaussian distribution of trajectory lengths. The central mark represented as a white dot indicates the median, and the gray line indicates the 25th and 75th percentiles. The whiskers extend to the most extreme data points not considered outliers. Trial corresponds to a stimulation pulse.

In summary, we observed a reduction of proprioceptive information during concurrent continuous stimulation of the cutaneous nerve also in the somatosensory cortex. This finding suggests that the effects that we observed in the spinal cord propagate through the higher layers of sensorimotor processing.

## Discussion

In this experimental study in four monkeys, we combined analysis of neural population dynamics with classical electrophysiology measures to analyze the impact of continuous electrical stimulation of the cutaneous afferents on the processing of proprioceptive information in the spinal cord. We found that the spinal sensorimotor computations of proprioceptive inputs were substantially disrupted when cutaneous afferents were concurrently stimulated and that this interference propagated to the brain. While limited to the stimulation of the cutaneous afferents, our findings suggest that artificially-generated neural input may disrupt ongoing neural processes that may be unrelated to the stimulation. More specifically, because of the highly shared neural architecture, even highly selective targeting of neural elements, like in our case the cutaneous afferents, can significantly undermine neural processing of seemingly unrelated neural functions, like proprioceptive percepts. Similar phenomena may occur in other regions of the nervous system and should therefore be studied. Hence, these results imply that efforts towards the development of naturalistic or biomimetic stimulation inputs should likely be employed in neurostimulation.

### Population analysis as a tool to explain network-level effects of electrical stimulationz

Electrical stimulation of the nervous system is widely applied in clinical practice and in clinical research trials in order to influence neural activity and ameliorate functions in a variety of disorders(*1, 2, 4, 6, 9*). The most overt applications are sensorimotor neuroprostheses where a clear relationship can be found between stimulation parameters and strength of elicited movements(*3, 4, 6, 43*) or evoked sensations(*1, 44–47*). For example, epidural spinal cord stimulation has been applied for both motor recovery as well as, more recently, for restoration of sensory feedback. We know that spinal cord stimulation recruits sensory afferents. In motor applications, the recruitment of large proprioceptive afferents leads to an increased excitability of spinal motoneurons, thereby promoting movement. Instead, in sensory applications, the recruitment of the same afferents should in principle produce controllable conscious sensory experiences, similar to those elicited by stimulation of the peripheral nerve. However, beyond this simplistic vision, there is a fundamental lack of knowledge into what happens to neural networks that receive inputs from these afferents. In fact, the highly shared neural infrastructures involve neural sub-networks that are meant to produce the desired function(*41, 48, 49*). For instance, the motor network produces movement, but other networks may be involved in other unrelated processes such as perception, error estimation during movement execution and autonomic function(*50*), among others(*48*). These additional sub-networks may also share inputs from the same afferents and would thus be perturbed by stimulation.

In our work, we constructed a toy model to study this specific problem in a controlled fashion as a proxy to understand, more generally, how artificial inputs can influence neural network function. We focused on the spinal network effects caused by stimulation of the cutaneous afferents on the neural processing of a proprioceptive pulse. This toy model exemplifies that an ongoing neural process (proprioceptive input processing) is perturbed when seemingly unrelated electrical stimuli application (inducing touch percepts) is applied with typical stimulation pattern (fixed 50Hz square pulses). To explore network effects, we used modern population analysis tools that enable the quantification of information processing in the spinal circuits via analysis of neural trajectories in the neural space(*33*). Specifically, we established a measure of proprioceptive information processing by quantifying neural trajectory lengths in spinal neural manifolds using intra-spinal population analysis(*36*). Through the quantification of the trajectory length, we assessed the effect of concurrent cutaneous stimulation on the spinal proprioceptive information. We showed the collapse of proprioceptive neural trajectories during concurrent stimulation of cutaneous afferents, in other words, a suppression of proprioceptive information processed in the spinal cord, according to our interpretation. Importantly, simple analysis of total spike counts showed that the results of our trajectory length quantification were not trivial. Indeed, averaged multiunit spiking activity in response to proprioceptive stimuli were expectedly the highest during concurrent stimulation of the cutaneous afferents. This is an obvious result as general spinal activity is increased by the 50 Hz artificial cutaneous inputs. Thus, the actual total neural activity in the spinal cord is higher during cutaneous stimulation. Yet, the population analysis allows to extract only activity that explains the variance generated by proprioceptive inputs processes, enabling to infer the proprioceptive components of the neural dynamics against the background of cutaneous activity. Hence, the use of neural manifolds for population activity enabled the quantification of the stimulation effects on these computations.

We validated results obtained with neural manifold analysis with classical electrophysiology inspecting peak-to-peak amplitude of afferent volleys, spinal cord grey matter response fields and multiunit activity. These measures indicated a reduction in proprioceptive information during concurrent cutaneous stimulation.

### Potential underlying neural mechanisms

While successful in visualizing network effects, population analysis cannot offer an explanatory value on the specific neural mechanisms responsible for this suppression. Pre-synaptic inhibition is a likely candidate(*51*). It is a well-known mechanism of sensory input gating that prevents transmission of excitatory post-synaptic potentials to neurons targeted by primary afferents(*39, 40*). In our experiments, the stimulation amplitude of the muscle nerve was the same across all conditions (i.e., fixed number of recruited afferents). Therefore, the reduction of unit responses to proprioceptive inputs during concurrent cutaneous afferent stimulation could be consistent with a reduction in synaptic inputs to these target units.

Nevertheless, the observed afferent volley reduction was not strong enough to explain complete diminishment of proprioceptive perception, which means that the disruption of neural trajectories was not caused only by reduced inputs, but also by affected processing. We refer to this other potential mechanism as the “busy line” effect. Continuous, non-natural stimulation of the cutaneous afferents may produce highly synchronized activity in spinal circuits, which may receive both proprioceptive and cutaneous inputs. However, when artificially synchronized cutaneous inputs reach the spinal cord, they may saturate these circuits and reduce their capacity to respond to additional inputs(*52*). When these neurons cannot be employed to process proprioceptive information, the neural network achieves a saturated state where no further processing can be carried out. This may explain why neural trajectories during cutaneous stimulation were displaced in the manifold space in a way that resembled a rigid geometric translation. Coincidentally, the modulated component of the neural dynamics was shorter or almost completely disrupted, suggesting that some of the neurons involved in performing the geometrical translation were not available to produce the modulated components of neural dynamics.

### Effects within the brain

We performed a large part of our analysis in the spinal cord, which is the first important layer of sensory processing, particularly, in regard to proprioception. However, conscious perception is processed at various layers above the spinal cord. Indeed, peak-to-peak amplitudes of cortex potentials evoked with muscle nerve stimulation were suppressed when overlapped with cutaneous input also in the sensory cortex area 2, which is known to integrate cutaneous and proprioceptive inputs(*48, 53*). Moreover, if cortical signals were independent from spinal and brainstem processes, when looking at the global cortical spike counts we would have expected higher spike counts during high-amplitude stimulation of the cutaneous nerve and lower spiking activity during low amplitude stimulation of the cutaneous nerve. Instead, we found higher or similar spiking activity when we used low-amplitude stimulation of the cutaneous nerve. This may be indicative of the fact that high amplitude stimulation may convey more cutaneous input but less proprioceptive input to the cortex because of sub-cortical cancellation(*39*). In contrast, cutaneous stimulation at a lower amplitude may mean less cutaneous input but more proprioceptive input to the cortex as a consequence of less cancellation occurring in sub-cortical structures. Nevertheless, these mechanistic conjunctures are strongly contingent on our experimental design. Future directions ought to design alternative experimental paradigms (i.e., including histological analysis) that uncover the spinal interneuron circuitry involved in the processing of proprioceptive information in response to concurrent input.

These overall results support the conclusion that conscious perception of proprioception may be also altered by sub-cortical interference. While this hypothesis cannot be tested in subjects with amputation because of their limb loss, recent data in humans with sensory incomplete spinal cord injury shows that spinal cord stimulation, which also recruits sensory afferents(*54*), reduces proprioception acuity during supra-threshold stimulation(*6*). This result in humans further supports our hypothesis and we believe that it demands further investigation.

### Conclusions and relevance for other brain circuits

Our results showed that electrical stimulation of sensory afferents can alter the processing of proprioceptive information within spinal circuits. Similar phenomena may occur in brain networks during deep brain stimulation, where similar continuous electrical pulses are delivered to thalamocortical projections and other large brain networks. In the brain, these effects, which in the spinal cord indicate the impossibility to appropriately process proprioception, could potentially alter cognitive processes unrelated to the stimulation goals within the cortex. A potential approach to minimize the interference of stimulation with ongoing neural processes is the use of “bio-mimetic” and model-based stimulation patterns (*23, 45, 55–57*). Instead of delivering unstructured and synchronized neural activity, they could produce more naturalistic patterns, thereby potentially avoiding these side effects. In conclusion, future stimulation strategies designs should consider the use of neural population analysis in order to analyze the effects of particular stimulation patterns on apparently unrelated neural network processes.

## Materials and Methods

### Animals

The study was conducted according to the guidelines of the Declaration of Helsinki, and approved by the local (Research Institute of Medical Primatology) Institutional Ethics Committee (protocol № 38/1, October 31, 2019) and by the University of Pittsburgh Animal Research Protections and IACUC (ISOOO17081).

Three adult Macaca Fascicularis and one Macaca Mulatta monkeys were involved in the study (MK1 - MK 42286, male, 4 years old, 3.5 kg, MK2 - MK 42588, male, 4 years old, 3.35 kg, MK4 - MK 42328, male, 4 years old, 3.48 kg; MK3 – 219-21, male, 7 years old, 11.5 kg). Data for all Macaca Fascicularis monkeys were acquired in the National Research Centre “Kurchatov Institute”, Research Institute of Medical Primatology, Sochi, Russia. Data for Macaca Mulatta monkey was acquired in the University of Pittsburgh, PA, US.

### Surgical procedures

All the surgical procedures were performed under full anesthesia induced with ketamine (10 mg/kg, i.m.) and maintained under continuous intravenous infusion of propofol (1% solution in 20 ml Propofol/20 ml Ringer 1.8 to 6 ml/kg/h), in addition to fentanyl (6-42 mcg/kg/hour) for the Macaca Mulatta, using standard techniques. Throughout the procedures, the veterinary team continuously monitored the animal’s heart rate, respiratory rate, oxygen saturation level and temperature. Surgical implantations were performed during a single operation lasting approximately 8 hours. We fixed monkeys’ heads in a stereotaxic frame securing the cervical spine in a prone and flat position. First, we implanted two silicon cuff electrodes (Microprobes for Life Science, Gaithersburg, MD 20879, U.S.A. and Micro-Leads, Somerville, MA 02144, U.S.A.) on the distal ends of the superficial branch and deep branch of radial nerve that we determined via anatomical landmarks. We then inserted EMG electrodes in the Extensor Digit. Communis, the Flexor Carpi Radialis and the Flexor Digit. Superficialis. We stimulated electrically two branches of the radial nerve and looked at the EMG response to verify which branch was the muscle branch and which one was the cutaneous branch. Second, we implanted the brain array using a pneumatic insertion system (Blackrock Microsystem). We performed a craniotomy and we incised the dura in order to get clear access to the central sulcus. We identified motor and sensory brain areas through anatomical landmarks and intra-surgical micro-stimulation. Specifically, we verified that electrical stimulation of the motor cortex induced motor responses in the hand muscles (**Fig. S1a**). We then determined the position of the somatosensory area S1 in relation to this spot and implanted the UTAH array electrode (Blackrock Microsystems, Salt Lake City, UT, U.S.A.) across Areas 1 and 2 (and Areas 3 and 4 for the Mulatta monkey), 1.2 mm lateral to midline and 3.1 mm deep using a pneumatic inserter (Blackrock Microsystems, Salt Lake City, UT, U.S.A.).

Finally, we performed a laminectomy from C3 to T1 and then directly exposed the cervical spinal cord. We implanted a 32-channel linear probe (linear Probe with Omnetics Connector 32 pins - A1x32-15mm-50-177-CM32; NeuroNexus, Ann Arbor, MI, U.S.A.) and a 64-channel linear probe (double linear Probe with Omnetics Connector 64 pins - A2x32-15mm-100-200-177; NeuroNexus, Ann Arbor, MI, U.S.A.) in the gray matter at the C5 spinal segment. To implant the probe, we opened the dura mater and created a small hole in the pia using a surgical needle through which penetration of the probe with micromanipulators was possible. We implanted the arrays using MM-3 micromanipulators (Narishige, Tokyo, Japan; David Koff Instruments for the Mulatta monkey). Experiments in all four monkeys were terminal. At the end the animals were euthanized with a single injection of penthobarbital (60 mg/kg) and perfused with PFA for further tissue processing.

### Electrophysiology in sedated monkeys

Monkeys were sedated with a continuous intravenous infusion of propofol that minimizes effects on spinal cord stimulation(*58*).

### Data analysis

We applied all data analysis techniques offline.

#### Pre-processing

We filtered raw signals recorded with 32 - electrode array implanted in the spinal cord, as well as signals documented with UTAH array in somatosensory cortex with comb filter to remove artefacts on 50 Hz/60 Hz (depending on the country where the experiments have been done) and its harmonics. We designed a digital infinite impulse response filter as a group of notch filters that are evenly spaced at exactly 50 Hz/60 Hz.

We detected single pulses of the deep branch of the radial nerve and extracted 430 ms of the inta-spinal and intra-cortical signal post stimulation.

#### Identification of sensory volleys resulting from muscle nerve stimulation

We were able to detect afferent volleys and the resulting gray matter response field evoked with muscle nerve stimulation in the spinal cord. We applied a 3rd order Butterworth digital filter and extracted the signal from 10 – 1000 Hz. Afferent volley is defined as a first volley after the stimulation pulse, occurring 3 - 4 ms after the stimulation (unique physiology of a single animal causes these variations) and followed with gray matter response field. We quantified the amount of processed proprioceptive information by neural network by measuring peak-to-peak amplitude values of the gray matter response field.

We applied a similar procedure to extract the muscle nerve evoked potentials recorded in the somatosensory cortex.

#### Characterization and quantification of neural spiking activity

We extracted neural spiking activity by applying a 3rd order Butterworth digital filter to the raw signal, separating the signal in frequency range from 800 Hz to 5000 Hz. We detected the spikes using thresholding algorithm(*59*). We determined the threshold value separately for each recording channel. To detect the accurate threshold value, we concatenated all data sets that we aim to analyze in a single file. All analyzed data sets were concatenated in a single file in order to detect proper threshold values. The same procedure was applied to intra-spinal and intra-cortical recordings.

Multiunit activity is presented in form of rasterplot and quantified with peri-stimulus time histogram (PSTH). Each dot in rasterplot represents a single detected spike. Every rasteplot row corresponds to the intra-spinal or intra-cortical activity perturbed with a single muscle nerve stimulus pulse. PSTH is quantified with mean event rate, defined as the average number of spikes across all single pulses of muscle nerve stimulation, within defined time frame.

### Neural manifold and trajectory length

To project the trajectories in the neural manifold, we previously computed multiunit spiking activity for each condition. We calculated the spiking activity for every 100 ms with a sliding window of 10 ms over 430 ms around each muscle stimulation pulse. We zero-padded the first repetition for 90 ms and then overlapped 90ms from the previous repetition for the rest of repetitions. The final step to smooth the spiking activity was the application of a Gaussian kernel (s.d. 20 ms) to the binned square-root-transformed firings (10 ms bin size) of each recorded multiunit. For each condition, this resulted in a matrix of dimensions C x T, where C is the number of channels in the dorso-ventral linear probe and T is the number of 10 ms windows in a repetition concatenated for all the repetitions within a condition. Subsequently, we proceed to eliminate noisy repetitions. We discarded those repetitions within each condition whose s.d. was greater than twice the total s.d. across all repetitions plus the total mean of the s.d. across all repetitions for that condition. For cortical data, we previously converted the distribution of s.d. to a lognormal distribution to apply this outlier cleaning rule.

To calculate the latent dynamics for each monkey, we z-scored each condition’s spiking activity before applying dimensionality reduction principal component analysis (PCA) to the concatenated spike counts. We selected the first 3 principal components that explained most of the variance (~65% for all 3 monkeys, 54% for one monkey) as neural modes to define the neural manifold. Convergence points were reached at the first 3 to 5 dimensions according to the eigenspectrum of each monkey. In this low dimensionality space, we proceeded by eliminating repetitions as a function of the distance to the median trajectory. In particular, we computed the median trajectory for each 10 ms window for each condition. For each window, we calculated the distance between the median trajectory and the trajectory elicited by each repetition within a condition. 25th and 75th percentiles of the obtained distances allowed to discard trajectories whose distance was greater than the 75th percentile plus 1.5 times the interquartile range of the averaged trajectory for that repetition across all 10 ms windows. The same criterion was applied for the lower range. Finally, we quantified the trajectory length for the remaining repetitions for each condition and calculated the average trajectory length across all 10 ms windows.

### Statistical Analysis

Multi-group significance comparison of data obtained from the neural manifold for each condition in all four monkeys was tested using Kruskal-Wallis test. The level of significance was set at ***p<0.005.

Significance of suppressed peak-to-peak amplitude values of afferent volleys was analyzed with one-way analysis of variance revealed (ANOVA). Each point represents the peak-to-peak amplitude as a response to a single stimulus pulse. Boxplots show: the central mark indicates the median, and the bottom and top edges of the box indicate the 25th and 75th percentiles, respectively. The whiskers extend to the most extreme data points not considered outliers, and the outliers are plotted individually using the ‘o’ symbol. The level of significance was set at ***p<0.001, **p<0.01 and *p<0.05.

## Acknowledgments

Data from MK1, MK2 and MK4 were acquired in Sochi in 2019, MK3 was acquired at the University of Pittsburgh in 2022. All data analysis was performed at the University of Pittsburgh and between University of Belgrade and ETH Zürich. The authors would like to thank Sara Conti for her help during pilot experimental procedures and Isabella Bushko for the design of the monkey figure.

## Author contributions

Study design: MC, SR

Designing and performing the surgical procedures: PM, NP, DK-O, DB, SO, DS, JG-M, PG, MC, SR

Performing all the experiments: all the authors

Data analysis and figure making: NK, JMB

Secured funding: MC, SR, PM, DS

Supervision: MC, SR

Writing—original draft: NK, JMB, SR, MC

Writing—review & editing: all the authors

## Data and materials availability

All data and code will be available upon reasonable request to the corresponding author.

## Supplementary Materials

**Fig. S1.**
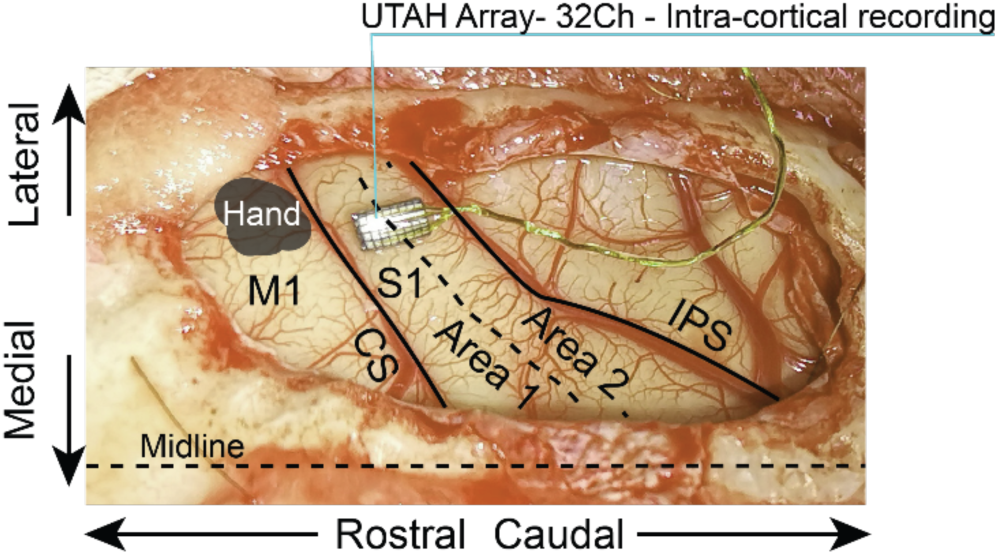
Experimental procedure and electrophysiology details. a) Representative picture showing the position of the UTAH array in relation to brain areas. We identified specific brain areas through anatomical landmarks and micro-stimulation of the cortex. We verified that a single pulse of stimulation delivered induced clear responses in the hand muscles. We determined the somatosensory area S1 in relation to the identified M1 anatomically and implanted the UTAH array electrode (Blackrock Microsystems) across Areas 1 and 2.

**Fig. S2.**
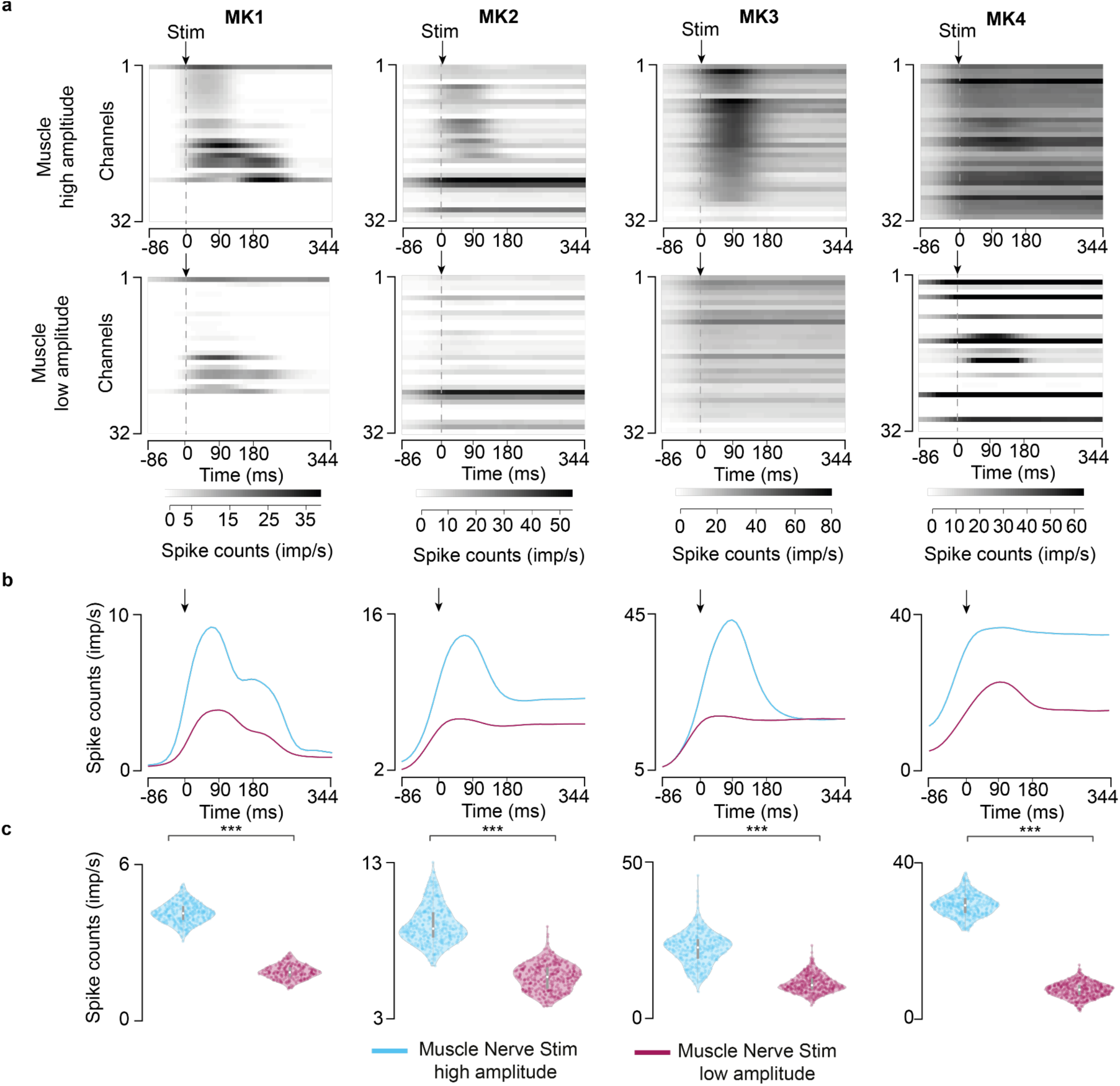
Spinal spiking activity induced by muscle nerve stimulation. a) Averaged spike counts for each channel. Spike counts were averaged across all trials for each stimulation condition for four monkeys. Averaged multiunit spike counts across all 32 channels, sorted by the highest spiking activity after the muscle nerve stimulation. b) Averaged spike counts. Spike counts were averaged across all trials and all channels for each stimulation condition for four monkeys. c) Statistical analysis of the spiking activity for each stimulation condition (***p<0.001; **p<0.01; *p<0.05; Kruskal-Wallis test with 387 and 234 points for muscle nerve stimulation at high amplitude and low amplitude, respectively, for MK1; 374 and 363 points, respectively, for MK2; 402 and 394 points, respectively, for MK3; 401, 353 and 343 points, respectively, for MK4). Violin plots: each dot corresponds to the computed trajectory length for a trial, forming a Gaussian distribution of trajectory lengths. The central mark represented as a white dot indicates the median, and the gray line indicates the 25th and 75th percentiles. The whiskers extend to the most extreme data points not considered outliers. Trial corresponds to a stimulation pulse.

**Fig. S3.**
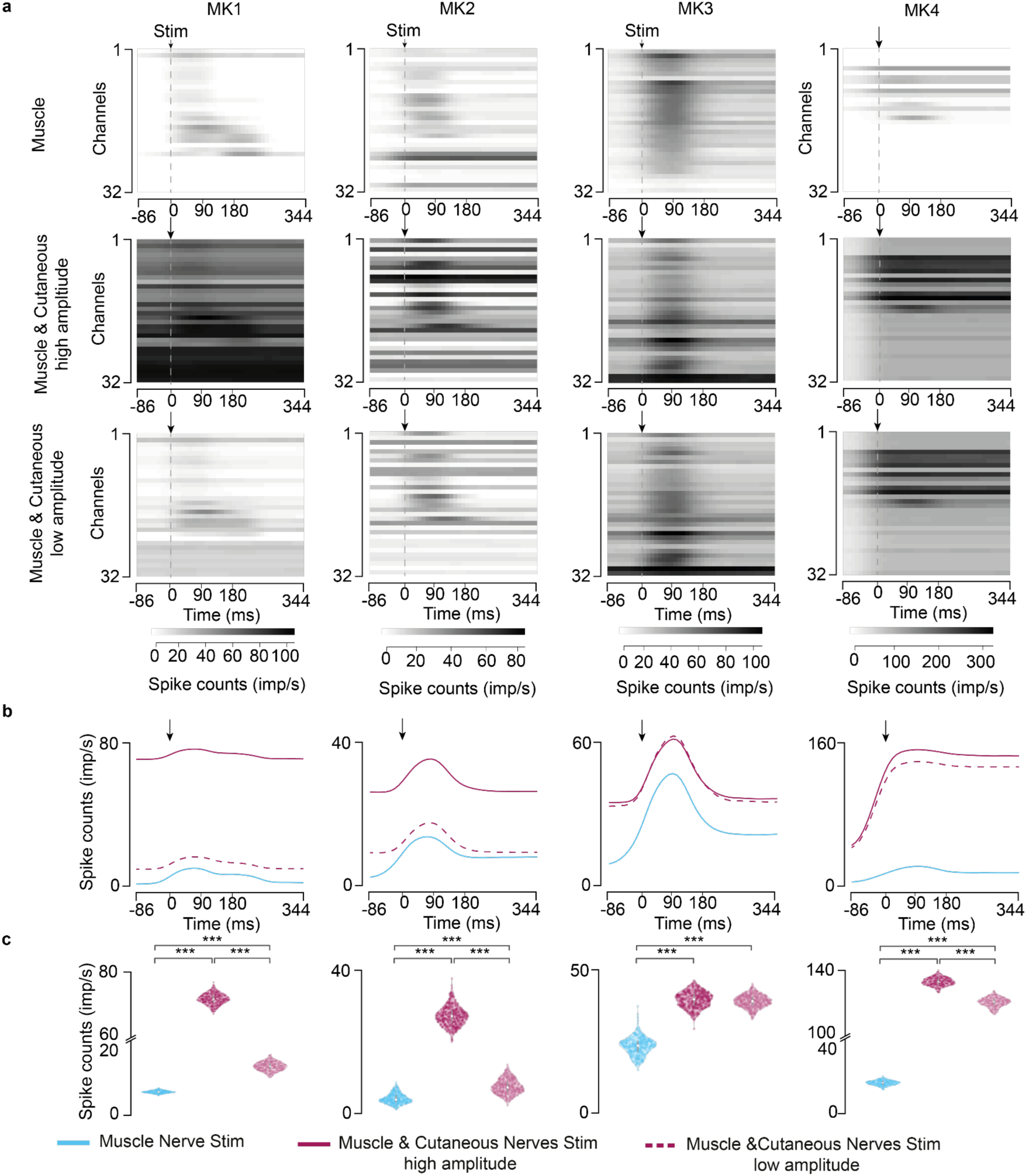
Spinal spiking activity induced by concurrent cutaneous nerve stimulation. a) Averaged spiking activity for each channel. Spike counts were averaged across all trials for each stimulation condition for four monkeys. Averaged multiunit spike counts across all 32 channels, sorted by the highest spiking activity after the muscle nerve stimulation. b) Averaged spiking activity. Spike counts were averaged across all trials and all channels for each stimulation condition for four monkeys. c) Statistical analysis of the spiking activity for each stimulation condition (***p<0.001; **p<0.01; *p<0.05; Kruskal-Wallis test with 387, 477 and 461 points for muscle nerve stimulation, concurrent cutaneous stimulation at high amplitude and low amplitude, respectively, for MK1; 374, 410 and 412 points, respectively, for MK2; 353, 400 and 399 points, respectively, for MK3; 401, 388 and 376 points, respectively, for MK4). Violin plots: each dot corresponds to the computed trajectory length for a trial, forming a Gaussian distribution of trajectory lengths. The central mark represented as a white dot indicates the median, and the gray line indicates the 25th and 75th percentiles. The whiskers extend to the most extreme data points not considered outliers. Trial corresponds to a stimulation pulse.

**Fig. S4.**
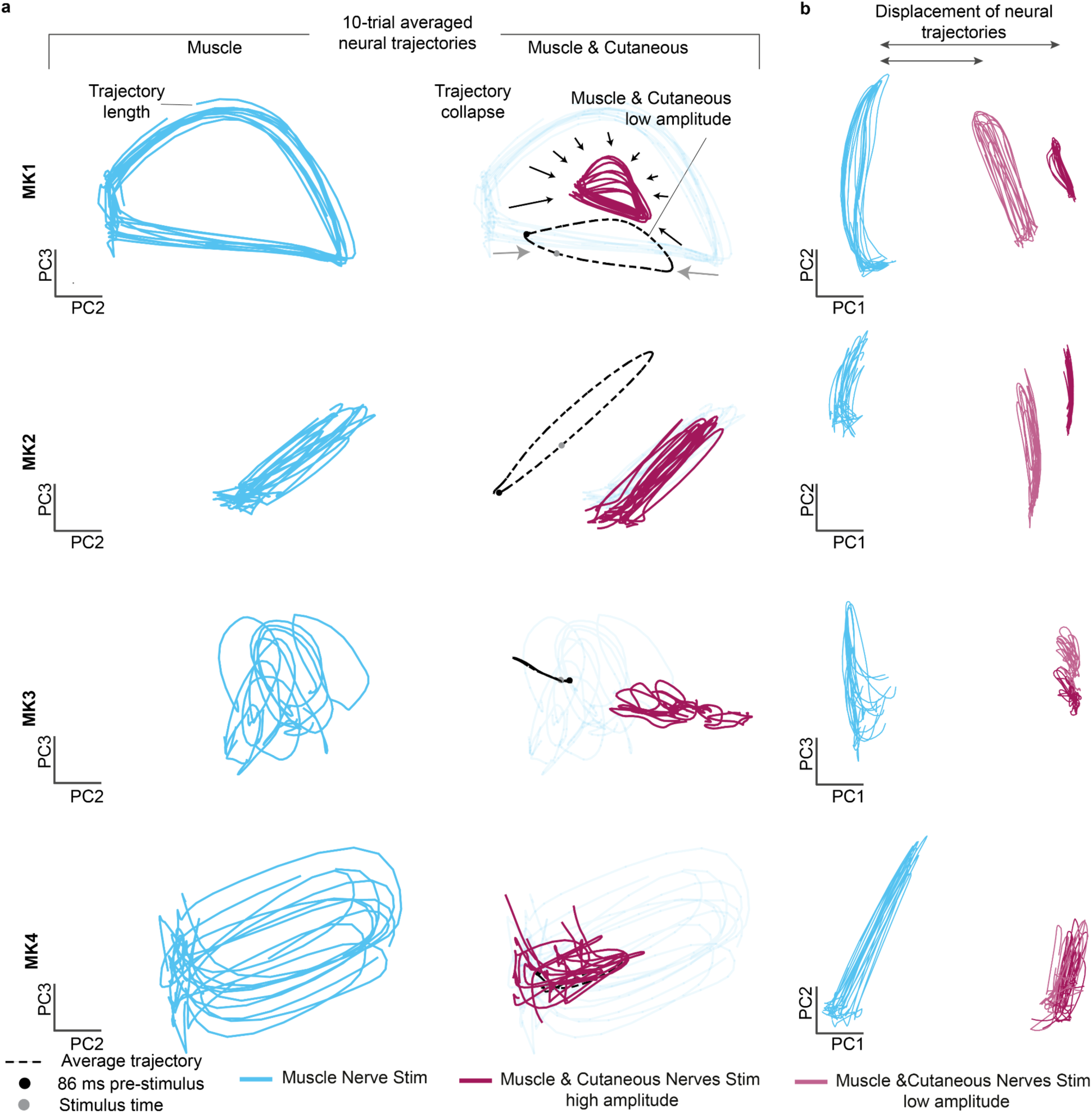
Intraspinal neural trajectories. a) Comparison of the neural trajectories induced by muscle nerve stimulation and concurrent cutaneous stimulation in all monkeys. Gray dashed lines indicate average trajectory for muscle and cutaneous nerves stimulation at a subthreshold amplitude. b) Visualization of the displacement of the neural trajectories across PC1 in all monkeys. The displacement is proportional to the spiking activity induced by each stimulation condition (i.e. the distance between neural trajectories induced by muscle nerve stimulation and concurrent cutaneous nerve at a low amplitude is lower than the distance between the neural trajectories induced by concurrent stimulation of the cutaneous nerve at a high amplitude).

**Fig. S5.**
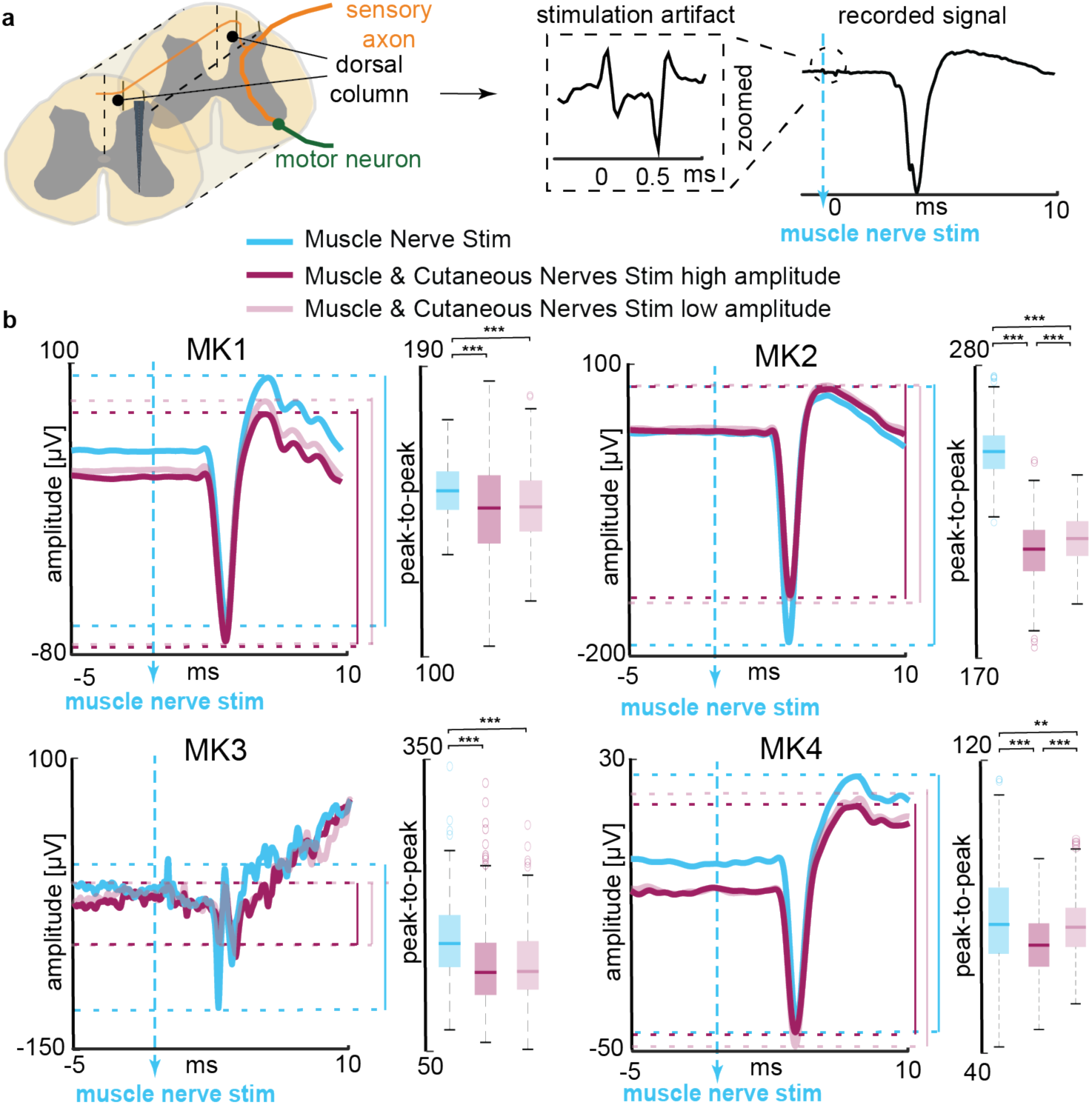
Proprioceptive afferent volley peak-to-peak amplitude suppression. a) Definition of the afferent volley. Triggered-average signal showed stimulation artifacts in the signal (zoomed insight) around the time of muscle nerve stimulation (pulse width: 0.5 ms) while the afferent volley appeared 3-4 ms after the stimulation (depending on the monkey). b) Afferent volleys in four monkeys. Afferent volleys as a response to proprioceptive nerve stimulation (cyan), with concurrent cutaneous nerve stimulation (magenta; high stimulation amplitude – solid color; low stimulation amplitude – semi-transparent). Volleys are given as an example of a single dorsal channel and are averaged across all muscle nerve stimulation pulses. We compared peak-to-peak amplitude values of afferent volleys over 2 conditions with one-way ANOVA with 300 points, where each point represents the peak-to-peak amplitude as a response to a single stimulus pulse. Boxplots: The central mark indicates the median, and the bottom and top edges of the box indicate the 25th and 75th percentiles, respectively. The whiskers extend to the most extreme data points not considered outliers, and the outliers are plotted individually using the ‘o’ symbol. Asterisks: ***p<0.001; **p<0.01.

